# Human emotional odours influence horses’ behaviour and physiology

**DOI:** 10.1101/2025.07.08.663503

**Authors:** Plotine Jardat, Alexandra Destrez, Fabrice Damon, Noa Tanguy-Guillo, Anne-Lyse Lainé, Céline Parias, Fabrice Reigner, Vitor H.B. Ferreira, Ludovic Calandreau, Léa Lansade

**Affiliations:** Institut Français du Cheval et de l’Equitation, Pôle développement, innovation et recherche, Nouzilly - France; INRAE, CNRS, Université de Tours, PRC, 37380, Nouzilly, France; Development of Olfactory Communication and Cognition Laboratory, Centre des Sciences du Goût et de l’Alimentation, Institut Agro Dijon, CNRS, Université de Bourgogne-Franche-Comté, Inrae, Dijon, France; UEPAO, INRAE, F-37380 Nouzilly, France

**Keywords:** human-animal relationship, equine cognition, *Equus caballus*, social cognition, semiochemicals, chemosignals

## Abstract

Olfaction is the most widespread sensory modality animals use to communicate, yet much remains to be discovered about its role. While most studies focused on intraspecific interactions and reproduction, new evidence suggests chemosignals may influence interspecific interactions and emotional communication. This study explores this possibility, investigating olfactory signals in emotional contagion from the example of human-horse interactions. Cotton pads carrying human odours from fear and joy contexts, or unused pads (control odour) were applied to 43 horses’ nostrils during fear tests (suddenness and novelty tests) and human interaction tests (grooming and approach tests). Principal component analysis showed that overall, when exposed to fear-related human odours, horses exhibited significantly heightened fear responses and reduced interaction with humans compared to joy-related and control odours. More precisely, when exposed to fear-related odours, horses touched the human less in the human approach test, gazed more at the novel object, and were more startled (startle intensity and maximum heart rate) by a sudden event. These results highlight the significance of chemosignals in interspecific interactions and provides insights into questions about the impact of domestication on emotional communication.

## Introduction

Olfaction, one of the most primitive senses, is probably the most widespread sensory modality used by animals to communicate. Evidence of communication through olfaction has been found across most animal groups [1], from insects to mammals, serving a wide range of biological functions. The most essential function is reproduction, where olfaction plays a crucial role, from sexual attraction (e.g., in snakes [2]) and mate recognition (e.g. in amphibians [3]) to mother-offspring recognition (e.g., in degu [4] or in cats [5]), highlighting the importance of chemical communication for fitness and survival. Olfaction is also involved in resource finding, with pheromone trails produced by ants to guide conspecifics toward food resources [6], and in spatial orientation, with wolf signalling biological fields through urinary marking [7]. Olfaction could also be involved in emotional communication, as odorous compounds are secreted in situations of danger, which are often associated with fear. For example, mustelids secrete sulphurous compounds when encountering predators [8]. Similarly, odours produced by decaying sea lampreys are avoided by other sea lampreys [9], which could be explained by fear being triggered in the recipients of this smell leading to avoidance of the related danger. This transmission of emotional information between conspecifics through olfaction has been explored specifically in a few studies. In cows and pigs, increased levels of stress (cortisol response, latency to feed) were observed when animals were exposed to urine from stressed conspecifics [10,11].

In humans, recent research into olfactory communication of emotions has highlighted the role of sweat produced by the apocrine glands in the armpits as carriers of emotional information [12,13]. Compounds present in sweat, such as adrenaline or androstadienone, have been suggested as potential carriers of this emotional information [14]. More than forty studies have demonstrated that such odours can influence the emotional state of the recipient [12]. For instance, chemosignals associated with fear or joy have been found to elicit corresponding fearful or joyful facial expressions in individuals exposed to them [14,15].

Most studies exploring the role of olfaction in social interactions have focused on intraspecific interactions. However, recent studies suggest that olfaction could play a role in interspecific emotional contexts as well. For example, dogs seem to perceive and react to human odours related to stress [16] or fear [17] and they show fewer interactions with a stranger and higher heart rates after smelling human odours related to fear than odours related to joy [18]. Mice and cows also appear to discriminate human odours associated with different emotional states [19], but to date the role of olfaction in interspecific interactions remains insufficiently understood. Further studies on domestic animals other than dogs would allow to better understand the phylogenetic extent of these phenomena and the impact of domestication on cognitive capacities.

Therefore, the aim of the present study was to further explore the role of emotional olfactory signals in interspecific interactions, and more precisely in human-animal interactions. Specifically, we studied horses and their reaction to human emotional odours. Indeed, horses have a presumably developed sense of smell [20] characterized by a convoluted olfactory bulb [21], and olfaction seems important for social interactions among horses [22,23]. Moreover, horses possess advanced cognitive skills related to humans. For example, they can recognize individual human faces in pictures, remembering them for at least six months [24,25], and they appear to detect our intentions [26]. They also perceive and interpret human emotions, reacting negatively to human voices and faces of anger compared to joy [27,28], and adapting their behaviour toward a human according to emotional cues they previously perceived [29]. They also perform social referencing based on human emotional cues [30], and they show cognitive integration of human emotional signals from different modalities [31– 33]. Moreover, recent studies revealed that horses can discriminate human body odours from contexts of fear and joy [34,35] and that they may perceive human joy odours as positive [35].

In this study, we tested whether horses’ behaviour and physiology can be influenced when they are exposed to odours emitted by humans feeling fear or joy. We first collected human armpit odours on cotton pads in contexts of joy and fear (volunteers watching horror and joyful videos). A control odour was also included, consisting in unused cotton pads. Then we conducted a series of tests with horses to evaluate whether these odours, when applied to the nostrils, modulate the behaviour and/or the physiology of horses. We used a protocol previously employed to assess the impact of odours (essential oils) on fear responses (novel object test and suddenness test) and on reactions toward humans (grooming test and human approach test [36–39]). Our hypothesis was that horses’ behaviour and physiology would vary according to the odour they were exposed to. Specifically, we expected horses to show higher fear responses and to be less prone to interact with humans when smelling the odour from the fear context compared to the control odour and the odour from joy context, and to show lower fear responses and be more prone to interact with humans when smelling the odour from the joy context, compared to the control odour and the odour from the fear context.

## Methods

### Ethics statement

This study was reported in accordance with ARRIVE guidelines [40]. It was approved by the Val de Loire Ethical Committee (CEEA VdL, Nouzilly, France, authorization number CE19—2022-1511-2). Animal care and experimental treatments complied with the French and European guidelines for the housing and care of animals used for scientific purposes (European Union Directive 2010/63/EU) and were performed under authorization and supervision of official veterinary services (agreement number F371752 delivered to the UEPAO animal facility by the veterinary service of the Département d’Indre et Loire, France). The horses were not food deprived during the experiment and did not undergo any invasive procedures.

Human participation in the experiment was carried out according to the Declaration of Helsinki and was approved by the Institutional Review Board of the University of Tours (authorization number 2022-029). All participants were fully informed about the general aims and methods of the study, and they provided written informed consent for the collection of samples as well as their use in the experiment. Data and code produced and used in this experiment are available in the following repository: https://doi.org/10.57745/T2RTDR

### Human odour collection

Human odours corresponding to fear and joy contexts were collected by a method identical to that described in [35]. Human axillary sweat odour was collected from 30 adult participants who volunteered to take part in the experiment. They were asked to follow dietary restrictions and use hygiene products provided by the experimenters (identical to [35]). For two days before sweat collection, they had to abstain from consuming certain produces known to affect body odours (chili pepper, spices, blue cheese, onion, garlic, cabbage, tobacco, and alcohol), refrain from using deodorant, perfume or scented lotion, and to bathe with a perfume-free soap that the experimenters provided. The morning before sweat collection, they had to rinse their armpits with clear water only.

Each participant watched two different videos on two different days (in a counterbalanced order), while wearing cotton pads under their armpits (7.5 × 7.5 cm, Euromedis, Neuilly-sous-Clermont, France) and a provided cotton t-shirt previously washed with clear water at 40°C. The video clip selected for the fear condition was a 20-minute excerpt from the movie *Sinister* [41]. The clips selected for the joy condition were the sketch *Les inventions* by Florence Foresti (a famous French humourist), *Singing in the rain*’s dance scene, *Have you ever seen a giraffe with a necklace* (Chez Wam production), *The Willy Waller 2006* (tetesaclaques.tv), *Birds* (Pixar), and *We go together* from *Grease*; for a total of 20 minutes.

After each session, participants placed the cotton pads in airtight sealed bags, which were stored in a freezer at −70 °C until presentation to horses. Participants indicated in a questionnaire whether they had thoroughly followed the dietary and hygienic instructions, and rated on 7-point Likert scales their extent of feeling angry, fearful, happy, sad, disgusted, neutral, surprised, calm, and amused. Samples from participants who had not followed the dietary and hygienic instructions or who did not take part in both sessions were excluded. Among the remaining samples we selected the 14 participants who experienced the most fear in the fear condition and the most joy in the joy condition (see Table S1), to match the number of horses included in the study (see Animals below). Ratings were combined and averaged to create indicators of high arousal (angry, fearful, happy, disgusted, surprised, amused), low arousal (neutral, sad, calm), positive affect (happy, calm, amused), and negative affect (angry, fearful, sad, disgusted - [14]). These scores revealed notable distinctions between the contexts for the selected participants, primarily in positive and negative affect scores, with some marginal variations in low arousal scores as well (Table 1). At the time of odour collection, unused cotton pads were placed in airtight sealed bags inside a freezer at −70 °C to be used as control odours.

**Table 1:** Experience of participants during the fear and joy contexts, transformed into dimensions of core affect (median[Q1,Q3])

|  | High arousal | Low arousal | Positive affect | Negative affect |
| --- | --- | --- | --- | --- |
| Fear context | 2.83[2.50,3.25] | 2.33[2.00,2.67] | 2.00[1.42,2.25] | 2.50[2.25,2.75] |
| Joy context | 2.67[2.50,2.83] | 3.17[2.75,3.58] | 5.67[5.08,3.00] | 1.00[1.00,1.00] |

### Animals

The study involved 43 Welsh mares (*Equus caballus*) aged 7.9 ± 2.2 years (mean ± s.d.) reared and living at the Animal Physiology Experimental Unit PAO (UEPAO, 37380 Nouzilly, France, DOI: 10.15454/1.5573896321728955E12), INRAE. These mares lived in groups in outdoor paddocks. Fodder and water were available ad libitum. These horses are used only for research purposes and are handled daily by humans. Horses were randomly assigned to one of three experimental groups: Joy group (n=14), to be exposed to human odours from the joy context; Control group (n=15), to be exposed to control odours (unused cotton pads), or Fear group (n=14), to be exposed to human odours from the fear context.

## Procedure

### Odour apparatus

In the morning, the cotton pads that were to be used during the day were transferred from the −70 °C to a −20 °C freezer that was close to the experimental area. The pads were thawed at room temperature (≈15°C) in the airtight bags 10 minutes before being used. The apparatus consisted of a disposable lycra muzzle (designed for this experiment) in which two cotton pads were stapled so that they would be in front of the horse’s nostrils once the apparatus was in place (Figure 1). Horses could breathe normally through this muzzle (they did not show any change in respiratory rate or breathing movements). The cotton pads presented to a horse by this apparatus were either two unused cotton pads (Control group) or two cotton pads from a single participant in one of the emotional contexts (fear context for the Fear group, joy context for the Joy group). The muzzles (a new one for each horse) had been previously washed with clear water without detergent and were always manipulated wearing gloves.

**Figure 1:**
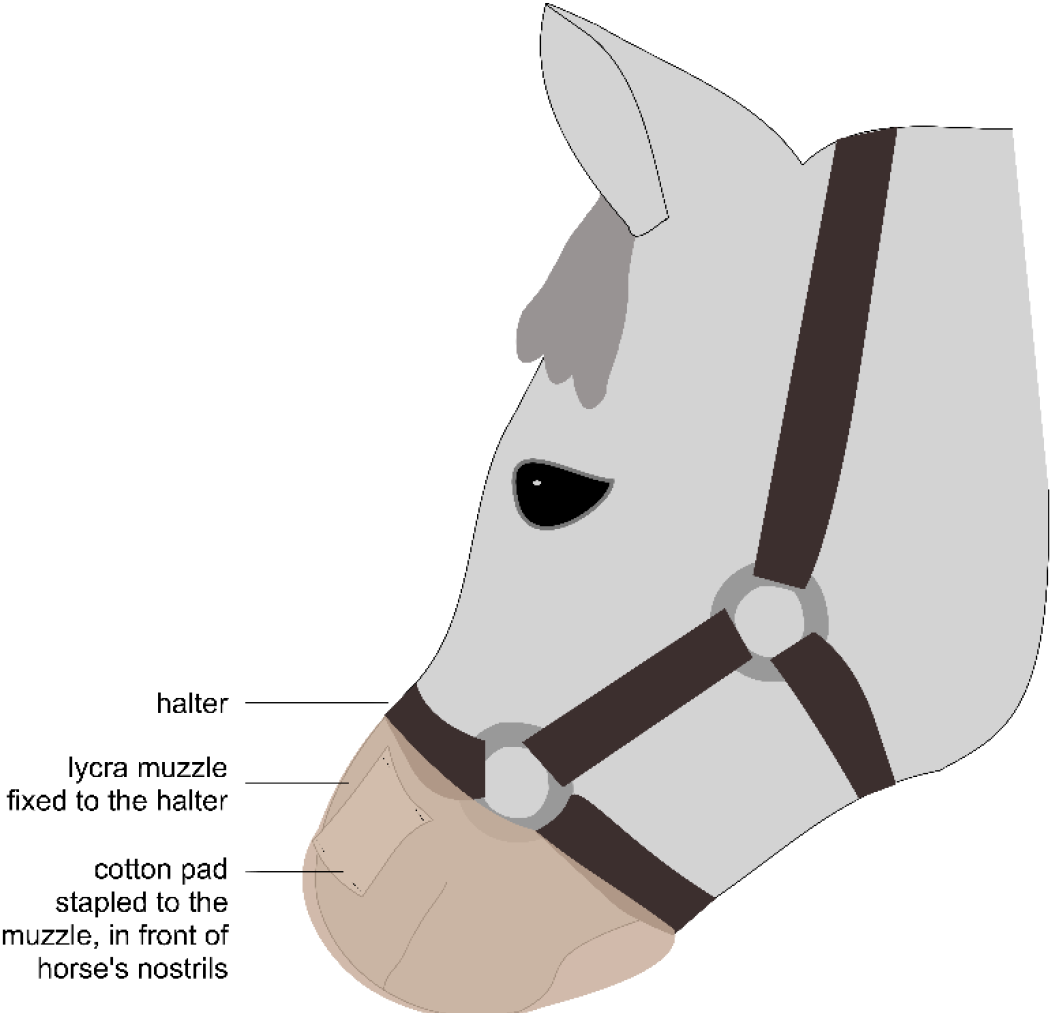
Muzzle used to expose horses to the odours. A lycra muzzle with cotton pads stapled in front of the horse’s nostrils was fixed to the halter.

### Behavioural tests

The tests took place in a 2.7 m x 8.1 m box designed to carry out reactivity tests for horses [36]. It was divided into 12 zones of equal surface, marked by tape (Figure 2). An audience horse from the tested horse’s herd was present at one end of the box, 1m from the fence, to avoid the tested horse to be stressed by a separation from conspecifics. She had been habituated to all the stimuli used in the tests and did not react to them. Four tests were conducted for each horse, two in the morning (morning series) and two in the afternoon (afternoon series), in a same order for all horses (from the less to the more frightening to the horse). For each test series, the cotton pads used were freshly thawed from the −20 °C freezer, with pads from the same donor used for a given horse in the morning and afternoon (each donor produced 4 pads, two of which were used for one horse in the morning and the other two for the same horse in the afternoon). On each testing day, the same number of horses were picked out randomly from each experimental group (Joy, Control or Fear) to be tested. Right before a series of tests began, the experimenter, who was familiar to the horses, led the horse from her paddock to the box, while two assistants prepared the odour apparatus (stapling the cottons pads to the muzzle) on a table a few meters from the test box. Only one of these assistants (hereby, the first assistant) was aware of which group the horse belonged to (Joy, Control or Fear), while the experimenter and second assistant were blind to the condition. Before entering the box, the muzzle with the odours was placed on the horse’s mouth and nostrils (Figure 1). The horse was then set free in the box for 1 minute (habituation phase). Then, the series of two tests were conducted. The second assistant, who was unaware of the horse’s group, recorded the horse’s behaviours live from outside the box, and the experiment was filmed by three cameras so that their notes could be completed afterwards if necessary. The morning series consisted of a grooming test and a free human approach test. The afternoon series consisted in a suddenness test and a novel object test.

**Figure 2:**
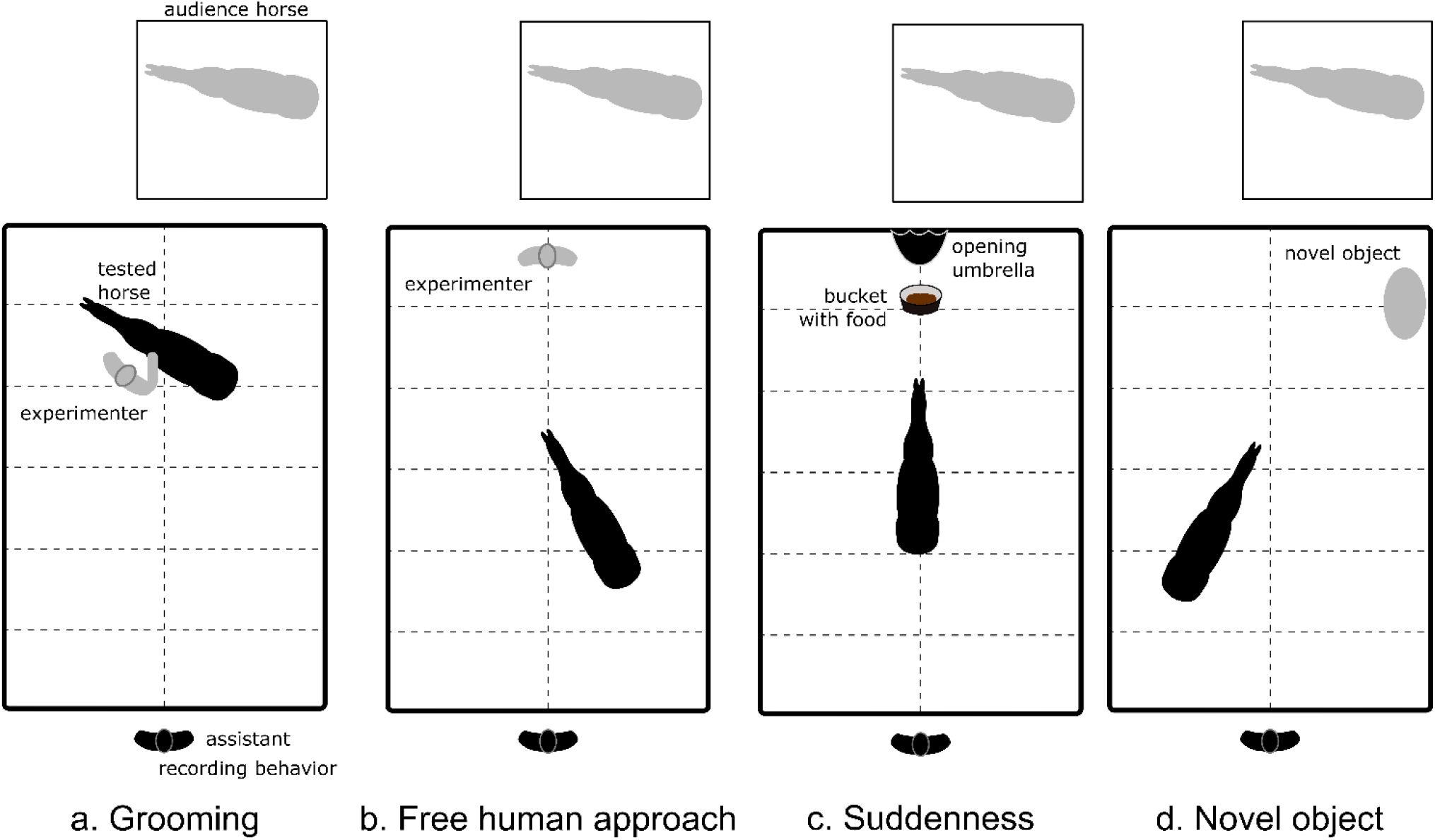
Behavioural tests. The experimental area was a 2.7 × 8.1 m box divided in 12 zones marked by tape. The audience horse was habituated to the stimuli, she could not touch or be touched by the tested horse. The tested horse was exposed to human odours from a joy or fear context or to control odours, according to her group.

#### Grooming test

The experimenter entered the box, tied a leading rope to the horse’s halter, and scratched her on the wither and neck in a standardized manner (see [42]) for two minutes (Figure 2a). The number of times the horse touched the experimenter was recorded, as a measure of her willingness to further interact with the experimenter.

#### Free human approach test

The experimenter set the horse free, walked to a predefined standing spot (Figure 2b), and stood for three minutes looking at her feet. The number of times the horse voluntarily touched the experimenter was recorded, as a measure of her willingness to interact with the experimenter.

#### Suddenness test

A hole was cut at the front of the muzzle so that the horse could eat (the cotton pads remained in place). A bucket with food pellets was placed on the ground on a predefined spot (Figure 2c). The first assistant placed a folded umbrella in a hole through the wall over the bucket, and waited for the horse to eat the food. Three seconds after the horse started eating, or after two minutes if the horse was still not eating from the bucket, the second assistant triggered the opening of the umbrella. The intensity of the startle response was recorded during the test as follows: 0 = no reaction, 0.025 = head raise, 0.25 = head raise and startle, 0.5 = step back, 1 = quarter turn (10°-90°, front legs <20cm above ground), 1.5 = violent quarter turn (10°-90°, front legs >20cm above ground) or half turn (90°-180°, front legs <20cm above ground), 2 = violent half turn (90°-180°, front legs >20cm above ground), 2.5 = very violent half turn (90°-180°, front legs > 40cm above ground, very rapid movement (<0.5s)), 3 = fall [43].

#### Novel object test

A novel item, unfamiliar to the horse, was placed inside the box on a predefined spot (Figure 2d). The object was a composition crafted from linoleum pieces and variously coloured and shaped plastic pieces connected by strings, its overall dimensions were 1mx1m. The horse was set free in the box for three minutes. The number of gazes at the object and the number of contacts the horse made with the object (touching the object with the nose or another body part) were recorded, as measures of her fear toward the object [37,43].

## Physiological measurements

Horses were habituated to the measurements from previous experiments.

### Salivary cortisol

Saliva samples were collected right before and right after each series of test (15 to 20 minutes apart) with Salivette® Cortisol (Sarstedt, Nümbrecht, Germany). Cotton buds were centrifuged at 3000×g for 20 min at 4 °C, and the saliva was stored at − 20 °C until analysis. The cortisol was measured in two 50 µL saliva aliquots per sample, using a competitive enzyme immunoassay in the INRAE Laboratory, Nouzilly, France. The assay sensitivity was 2ng/mL. For each sample the mean concentration of cortisol in the two aliquots was calculated. Then, for each horse the variation in salivary cortisol was calculated as the difference between the mean level before and after the test series.

### Heart rate

Horses were equipped with a heart rate monitor (Polar Equine RS800CX Science, Polar Oy, Finland), recording RR data during the human approach, suddenness, and novel object tests (in the grooming test, grooming movements on the horse’s wither prevented the monitor from functioning normally). RR data were then extracted from the recordings and converted to heart rate values. A visual inspection of this data was performed to detect artefactual beats (as recommended by [44]). Numerous artefacts were spotted, so data were cleared up based on the following algorithm (code used to apply these rules is available in the online repository, see Figure S1 and Table S2 for more details): for each data point, if it had a difference of more than 35 bpm with the previous point, it was considered artefactual and removed from the dataset. If for a horse more than 30% of points concerning a test were artefactual, these data were considered unreliable and all data concerning this test for this horse were removed from the dataset. This was repeated three times. Mean and maximum heart rate were then calculated for each test.

## Precautions to limit odour contaminations

The odour samples were always manipulated with disposable gloves and an FFP2 mask. In order to limit the spread of her own body odour, the experimenter took two showers a day (one before each testing half-day) using unscented soap, and wore a swimming cap during the tests. She and the two assistants did not wear any perfume or deodorant for the time of the experiment. She wore gloves, an FFP2 mask and clean overalls that were changed for each horse, and boots that were cleaned with clear water between each horse. The faeces were picked up and the box was thoroughly cleaned with a water jet after each horse finished the tests. The muzzle used for each horse was disposed of after each horse finished the tests.

## Statistical analysis

All statistical analysis were performed using R 4.3.1 [45], and figures were generated using the packages *ggplot2* [46] and *factoextra* [47]. The significance threshold was set at α ≤ 0.05 and tendencies were considered for p-values ≤ 0.1.

### Behavioural variables

To test whether horses reacted differently to the tests according to the odour they smelled, we explored each recorded behaviour with generalized linear mixed models (GLMM) from the package *glmmTMB* [48], using linear and Poisson distributions as appropriate for the given variable (see Table S3). For each variable, a model was constructed to assess the effect of the group (Joy, Control or Fear) on the behaviour, and compared to the null model via likelihood-ratio (chi-square) test (Table S3). Distributions, within-group variance and homoscedasticity of the residuals were checked using the package *DHARMa* [49]. An chi-square test (Type II Wald) was then performed on each model to assess the influence of the group on the response variable using the function *Anova()* from the package *car* [50]. Post hoc tests based on Tukey’s method were performed when necessary with the function *emmeans()* from the package *emmeans* [51].

To visualize overall differences between groups across behaviours, a principal component analysis (PCA) was performed using the function *prcomp()* from the *factoextra* package [47]. Correlations between all the variables were checked using the function *corrplot()* from the package *corrplot* [52] (Figure S2). To test whether overall differences in behaviour differed significantly between groups, we explored the first two components of the PCA with generalized linear mixed models (GLMM), following the same method as for individual variables (i.e., we tested whether the coordinates of individuals along these first two components differed according to the group).

### Physiological variables

The evolution of cortisol level (before/after each test series), and the mean and maximum heart rate were analysed using linear mixed models (LMM) from the package *glmmTMB* [48]. For each variable, a model was built to assess the effect of the group on the physiological parameter. For the evolution of cortisol level, the series (morning or afternoon) was added as a random effect, to assess for potential variability introduced by cortisol circadian rythms [53]. The models were compared to the null model via likelihood ratio (chi-square) test (Table S4). Distributions, within-group variance and homoscedasticity of the residuals were checked using the package *DHARMa* [49]. For models that were significantly better than the null model, an chi-square test (Type II Wald) was performed to assess the influence of the group on the response variable using the function *Anova()* from the package *car* [50]. Post hoc tests based on Tukey’s method were performed when necessary with the function *emmeans()* from the package *emmeans* [51].

## Results

Descriptive statistics and effect sizes are given for all variables in Table S5

### Behavioural variables

The number of touching the experimenter during the human approach test and across all tests, along with the startle intensity in the suddenness test and the number of gazes toward the novel object differed significantly according to the group (Table 2). Post-hoc tests revealed horses in the Fear group touched the experimenter less and showed higher fear responses compared to the Control and/or the Joy groups, according to the variable (Table 2, Figure 3). The number of touching the experimenter during the grooming test and the number of contacts with the novel object tended to differ significantly between groups (Table 2). Post-hoc tests revealed that horses in the Fear group tended to touch the experimenter less and to make less contacts with the novel object compared to the Joy group (Table 2, Figure 3).

**Table 2:**
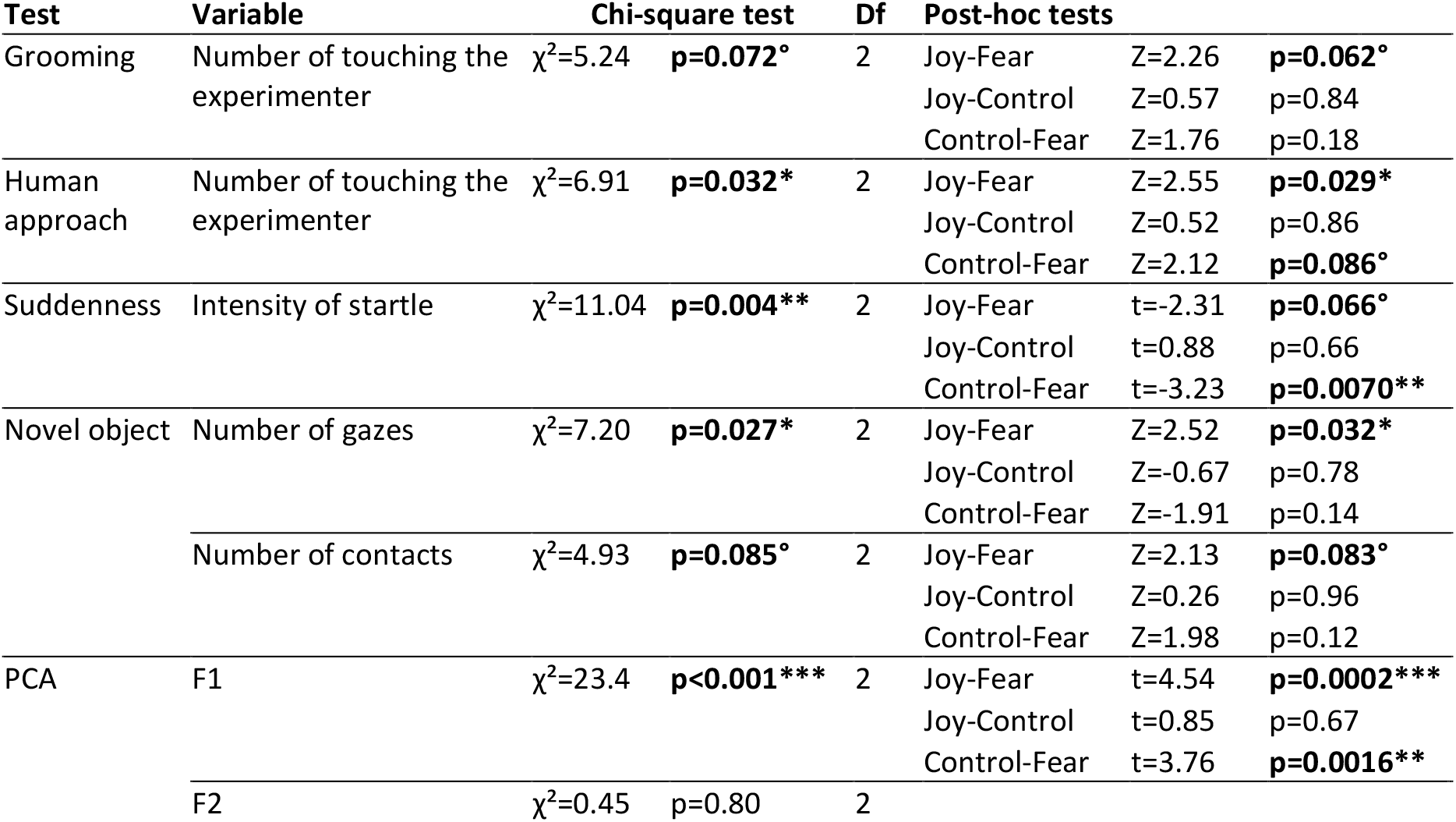
Results of the tests performed for each behavioural variable. (° p≤0.1, * p≤0.05, ** p≤0.01, *** p≤0.001). Joy: horses exposed to cotton pads from humans watching a joyful video, Control: horses exposed to unused cotton pads, Fear: horses exposed to cotton pads from humans watching a horror video.

**Figure 3:**
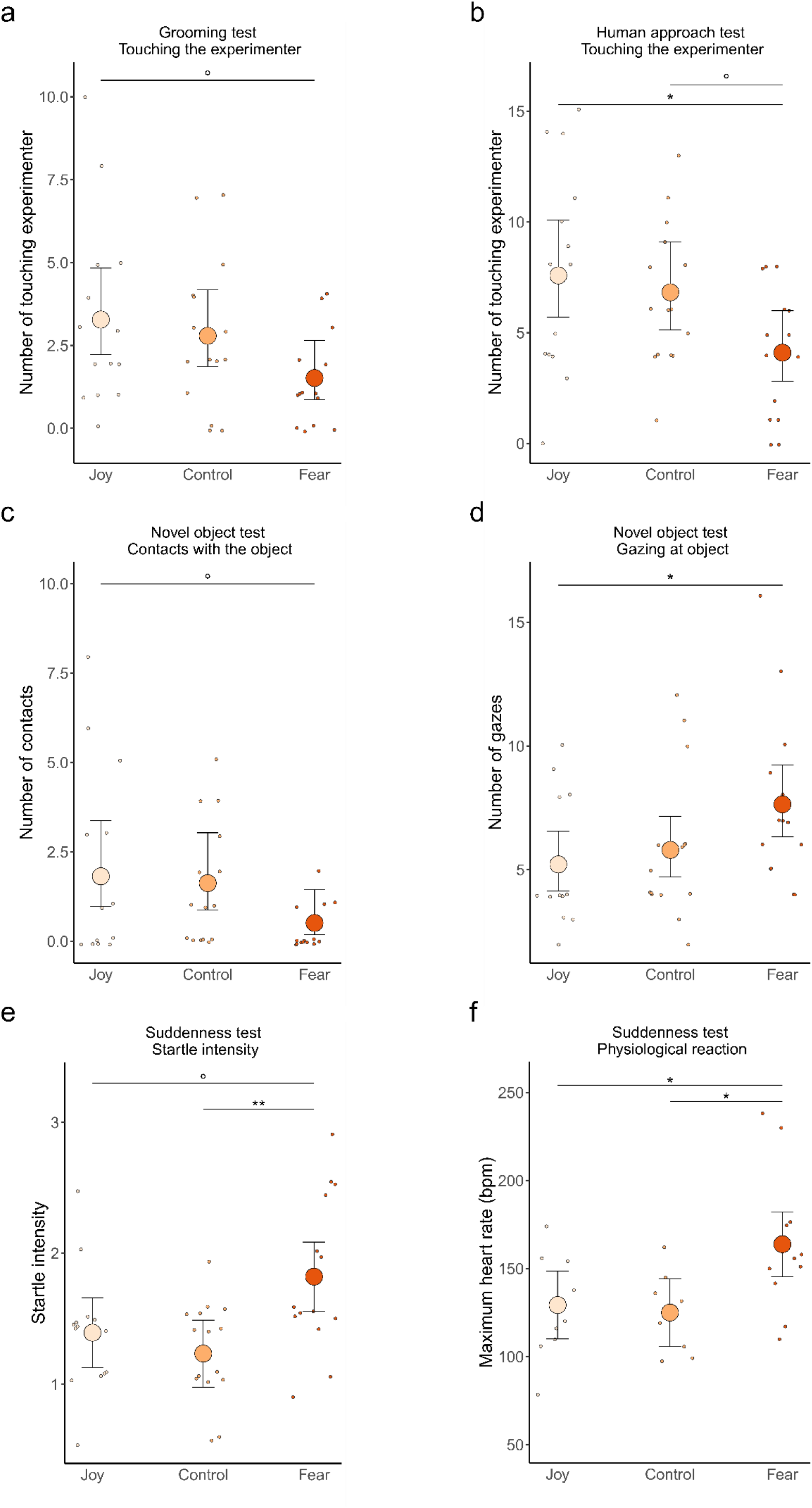
Behaviour and physiology of horses during the tests according to the odour smelled. a. Number of touching the experimenter in the Grooming test b. Number of touching the experimenter in the Free human approach test, c. Number of contacts with the objects in the Novel object test, d. Number of gazes at the object in the Novel object test, e. Startle intensity in the Suddenness test, f. Maximum heart rate in the Suddenness test. Joy: cotton pads from humans watching a joyful video, Control: unused cotton pads, Fear: cotton pads from humans watching a horror video. Individual data points with mean and standard error from the models presented in Tables S3 and S4. °p≤0.1, *p≤0.05, **p≤0.01.

The two first factors of the PCA explained 62.4% of the total variability (Table 3, Figure 4). Coordinates along the first factor differed significantly according to the group, post-hoc tests revealed horses in the Fear group had lower coordinates along this factor (they were more on the left of the graph) compared to the Joy and Control groups (Table 2). This factor (F1) corresponded to horses’ willingness to interact with the experimenter (positive values: willing to interact) and to the intensity of fear reactions (negative values: stronger fear reactions).

**Table 3:** Correlations between the variables included in the PCA and the first two factors.

| Test | Variable | F1 (37.1%) | F2 (25.3%) |
| --- | --- | --- | --- |
| Grooming | Number of touching the experimenter | 0.48 | -0.54 |
| Human approach | Number of touching the experimenter | 0.59 | -0.13 |
| Suddenness | Intensity of startle | -0.41 | -0.50 |
| Novel object | Number of gazes | -0.28 | 0.38 |
|  | Number of contacts | 0.42 | 0.54 |

**Figure 4:**
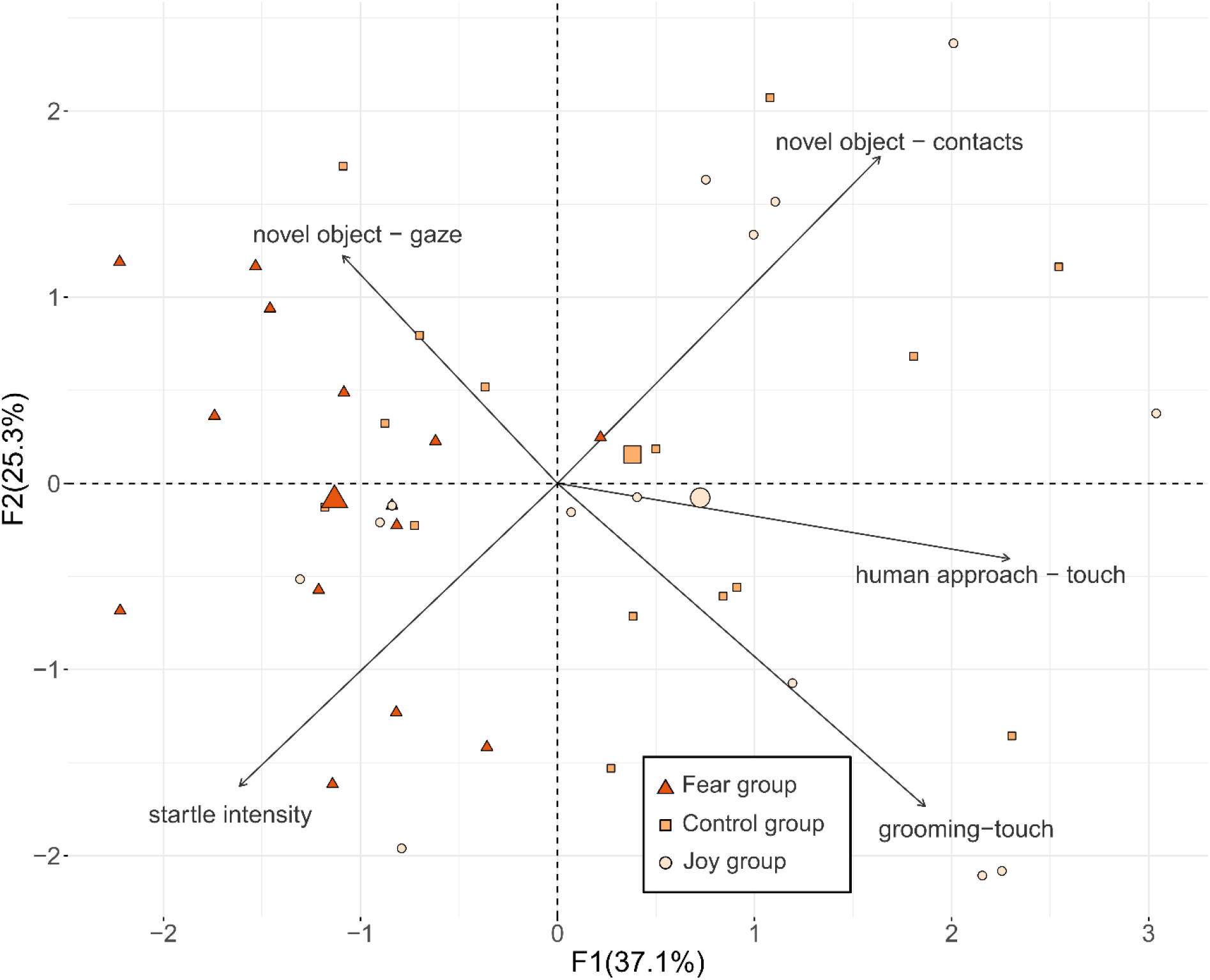
Horses’ behaviour during the tests as described by the PCA. Fear group: horses exposed to cotton pads from humans watching a horror video, Control group: horses exposed to unused cotton pads, Joy group: horses exposed to cotton pads from humans watching a joyful video. The large points represent the barycentre for each group and the small points individual data points from the PCA.

### Physiological variables

The maximum heart rate in the suddenness test differed significantly according to the group (Table 4). Post-hoc tests revealed that the maximum heart rate of horses in the Fear group was significantly higher than that of horses in the Joy group and in the Control group (Figure 3f, Joy-Fear: t=-2.66, p=0.033, Joy-Control: t=0.32, p=0.94, Fear-Control: t=-2.99, p=0.016). The other physiological parameters did not differ significantly according to the group (Table 4).

**Table 4:**
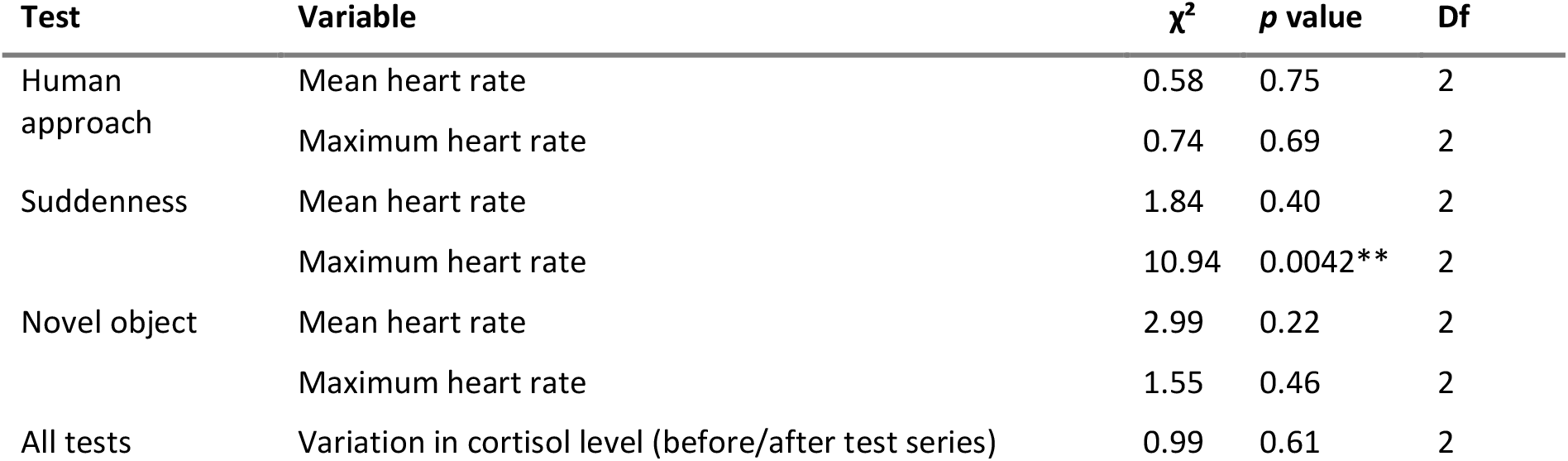
Results of the chi-square tests for each physiological variable. (** p≤0.01).

| Test | Variable | $\chi^2$ | $p$ value | Df |
| --- | --- | --- | --- | --- |
| Human approach | Mean heart rate | 0.58 | 0.75 | 2 |
|  | Maximum heart rate | 0.74 | 0.69 | 2 |
| Suddenness | Mean heart rate | 1.84 | 0.40 | 2 |
|  | Maximum heart rate | 10.94 | 0.0042** | 2 |
| Novel object | Mean heart rate | 2.99 | 0.22 | 2 |
|  | Maximum heart rate | 1.55 | 0.46 | 2 |
| All tests | Variation in cortisol level (before/after test series) | 0.99 | 0.61 | 2 |

## Discussion

In this study, we showed that horses’ behaviour and physiology were influenced when exposed to odours emitted by humans when feeling fear or joy. When exposed to the odour from the fear context, horses touched the experimenter less, gazed more at the novel object, and were more startled in the suddenness test, compared to odours from the joy context or control odour. These behaviours indicate higher fear responses [37,38,43] and less willingness to interact with humans [39] when exposed to fear-related odours. Principal component analysis confirmed that horses were overall more afraid and less willing to approach humans when exposed to fear context odours, as their coordinates on the first axis were significantly lower in the Fear group compared to the Joy and Control groups.

More than just a reaction to odours, there appears to be an analogy between the emotional state of the donor (human) and that of the receiver (horse), which could be the result of emotional contagion, a phenomenon defined as the transmission of an emotional states from an emitter to a receiver [54]. Indeed, the observed responses indicate a shift towards a more negative and higher arousal emotional state in horses, consistent with the emotional valence and arousal of the human emitters, thereby meeting criteria used to evaluate emotional contagion [55]. This suggests that emotional contagion of fear through odours, previously observed among humans [56–58], could also occur interspecifically. The reactions observed in horses could be spontaneous, deriving from autonomic pathways connecting olfactory receptors to emotional reactions and behaviours; or they could result from associations formed by horses between some odours and the situations in which they are usually present (for example, horses could have associated odours emitted by fearful humans to stressful contexts). Indeed, horses are known to associate some characteristics of human stimuli (like the nature of a voice) to the valence of co-occurring interactions [59].

The fact that different species appear to respond to each other’s emotional chemosignals is interesting, as emotions have previously been viewed as mainly internal states that serve to regulate individual behaviours [60,61] and produce signals that inform conspecifics on danger or resources [54,62]. The fact that these signals transcend species boundaries suggests that they could also play a role in interspecific interactions, especially between humans and domestic mammals. Fear derives from the perception of danger [63], and is probably the most widely shared emotion in the animal kingdom. Its function is to trigger a fight-or-flight response [63], so emotional contagion of fear as observed in this experiment could be adaptive for animals to warn each other of danger, even when it involves different species. Olfactory signals provoking emotional contagion of fear appear particularly effective as odours remain effective in darkness, in the presence of obstacles, and persist even after the emitter has departed [1].

To date, responses to interspecific emotional signals have been observed mainly in domestic animals. For example, horses but also dogs, cats, cattle and goats can discriminate human emotional expressions from visual, auditory or olfactory signals [64]. Goats [65], horses [32] and cats [66] show preferences for positive emotions over negative ones, and dogs [67], horses [31,32], and cats [68] may even form mental representations of human emotions. This raises the question of whether these capacities emerged from domestication. Indeed, domestication has led to the “domestication syndrome,” featuring traits like larger body size, increased reproduction, and reduced anti-predator behaviours [69,70], possibly including enhanced socio-cognitive capacities toward humans [71,72] and the ability to react to human emotional odours. However, domestic species do not necessarily surpass their wild counterparts in socio-cognitive abilities [70], potentially due to the ancestral selection of wild species based on these traits. Specifically, regarding olfaction, responses to human emotional odours have been observed primarily in domesticated animals (horses [34,35] and dogs [17,18,73,74]), which could be explained by their ancestral species already having a particular aptitude for this. However, it is also possible that mammals possess receptors inherited from a common ancestor that allow detection of common molecules produced in sweat across species. Future studies, including chemical analysis of emotional body odours, could determine if this capability is unique to domesticated animals and identify compounds that trigger fear to be tested in interspecific experiments on behavioural and physiological responses [75,76].

In this study, we did not find significant differences in horses’ behaviour between odours from the joy context and unused cotton pads. However, results indicate a gradient in horses’ reactions across the Joy, Control, and Fear groups, as depicted in graphical representations (Fig3a-d&f); and the detection of significant differences between the Fear group and the Joy group but not between the Fear group and the Control group suggests horses’ reaction to the control odours was not equivalent to that to the odour from the joy context. Future studies employing habituation-discrimination tests similar to those conducted previously [35] could further elucidate horses’ ability to differentiate between joy and control odours and explore their potential effects on horses’ emotional states. In humans, odours from a joy context have been found to induce a facial expression and a way of looking at images indicative of happiness in the receivers [14], suggesting that these odours differ in nature to control odours and could also influence the behaviour of individuals from other species. Although it is less evident than for fear, reacting to interspecific signals of positive emotions could be adaptive. For species that have regular interspecific interactions (like domestic animals or species involved in mutualistic relationships) it could help to promote social bonds [54]. Moreover, happiness is part of the reward acquisition system, of which the role is to promote reward-seeking behaviours like feeding [60], and emotional contagion of happiness could encourage resource identification.

Last, in this study no significant differences in cortisol levels were observed, which could be due to circadian rhythms inducing marked variability [53], or to this parameter varying over longer periods than our tests allowed (less than an hour per horse), so it would be interesting to repeat this investigation over more extended periods [77,78]. However, increased maximum heart rate during the suddenness test suggests physiological responses to interspecific fear-related odours do occur, which is in line with the function of emotions to recruit appropriate physiological resources in response to a stimulus, supporting survival [79].

## Conclusion

In this study, we showed that human emotional chemosignals influence horses’ behaviour and physiology, with human odours from a fear context inducing higher fear responses in horses and lower interactions with humans. This highlights the significance of chemosignals in interspecific interactions and provides new insights into questions about the impact of domestication on emotional communication.

## Acknowledgements

We thank the staff from UEPAO for technical help, and all human participants who watched the joy and fear videos and donated their sweat for the experiment.

## Declarations

### Author contributions

PJ : Formal Analysis, Funding acquisition, Investigation, Methodology, Software, Visualization, Project administration, Writing – original draft; AD: Conceptualization, Supervision, Methodology, Writing – review & editing; FD: Conceptualization, Supervision, Methodology, Writing –review & editing; NTG: Data curation, Formal analysis, Investigation, Methodology; AL: Methodology, Validation, Investigation; CP: Data curation, Investigation, Methodology; FR: Resources, Conceptualization, Methodology; VF: Conceptualization, Supervision, Methodology, Writing – review & editing; LC: Conceptualization, Supervision, Methodology, Writing – review & editing; LL: Conceptualization, Supervision, Methodology, Resources, Project administration, Writing –review & editing.

### Data availability

Data and code generated and analysed in this study have been deposited at https://doi.org/10.57745/T2RTDR and are publicly available as of the date of publication.

### Use of Artificial Intelligence (AI) and AI-assisted technologies

During the preparation of this work the authors used ChatGPT in order to improve readability and language. After using this tool, the authors reviewed and edited the content as needed and take full responsibility for the content of the publication.

