## supplementary information for "Human emotional odours influence horses’ behaviour and physiology"

**Table S1: Selection of the odour samples**

| Participant number | Followed hygienic and dietary instruction | Rating of feeling fearful during the Fear video | Rating of feeling joyful during the Joy video | Samples selected for experiment |
|---|---|---|---|---|
| 1 | yes | 2 | 7 | no |
| 2 | yes | 3 | 5 | no |
| 3 | yes | 3 | 7 | no |
| 4 | yes | 6 | 3 | yes |
| 5 | yes | 3 | 7 | no |
| 6 | yes | 4 | 6 | no |
| 7 | yes | 7 | 5 | yes |
| 8 | yes | 4 | 4 | no |
| 9 | yes | 2 | 5 | no |
| 10 | yes | 4 | 5 | no |
| 11 | yes | 6 | 5 | yes |
| 12 | no | 4 | 6 | no |
| 13 | yes | 6 | 6 | yes |
| 14 | yes | 6 | 6 | yes |
| 15 | yes | 3 | 6 | no |
| 16 | yes | 5 | 6 | yes |
| 17 | yes | 3 | 3 | no |
| 18 | yes | 5 | 5 | yes |
| 19 | yes | 5 | 5 | yes |
| 20 | no | 1 | 5 | no |
| 21 | yes | 1 | 4 | no |
| 22 | yes | 6 | 5 | yes |
| 23 | yes | 6 | 6 | yes |
| 24 | yes | 5 | 3 | yes |
| 25 | yes | 5 | 6 | yes |
| 26 | yes | 5 | 6 | yes |
| 27 | yes | 5 | 6 | yes |
| 28 | no | 4 | 5 | no |
| 29 | yes | 4 | 7 | no |
| 30 | yes | 4 | x | no |

Samples from participants who had followed the hygienic and dietary instructions were excluded. Among the remaining samples, those form the participants who had felt the most fearful during the fear video and joyful during the joy video were selected.

**<u>Table S2</u>: Number of horses whose heart rate data were included in the analysis for each test.** The initial group sizes were n=14 for the Joy group and the Fear group and n=15 for the Control group

| Test | Joy group | Control group | Fear group |
|---|---|---|---|
| Human approach | 13 | 13 | 13 |
| Suddenness | 10 | 10 | 11 |
| Novel object | 9 | 12 | 13 |

**<u>Table S3</u>: Models used for the analysis of behavioural variables and results of their comparison to the null model** (° p≤0.1, * p≤0.05, ** p≤0.01, *** p≤0.001)**.**

| **Test** | **Response variable (y)** | **Family** | **Model formula** | **χ²** | **DF** | ***p* value** |
|---|---|---|---|---|---|---|
| Grooming | Number of touching the experimenter | Generalized Poisson | y ~ Group | 5.59 | 2 | 0.061° |
| Human approach | Number of touching the experimenter | Generalized Poisson | y ~ Group | 7.27 | 2 | 0.026* |
| Suddenness | Intensity of startle | Gaussian | y ~ Group | 9.82 | 2 | 0.0074** |
| Novel object | Number of gazes | Poisson | y ~ Group | 7.09 | 2 | 0.029* |
| | Number of contacts | Generalized Poisson | y ~ Group | 6.11 | 2 | 0.047* |
| PCA | F1 | Gaussian | y ~ Group | 18.68 | 2 | <0.001*** |
| | F2 | Gaussian | y ~ Group | 0.38 | 2 | 0.83 |

**<u>Table S4</u>: Models used for the analysis of physiological variables and results of their comparison to the null model** (** p≤0.01)**.**

| **Test** | **Response variable (y)** | **Family** | **Model formula** | **χ²** | **DF** | ***p* value** |
|---|---|---|---|---|---|---|
| Human approach | Mean heart rate | Gaussian | y ~ Group | 0.57 | 2 | 0.75 |
| | Maximum heart rate | Gaussian | y ~ Group | 0.73 | 2 | 0.69 |
| Suddenness | Mean heart rate | Gaussian | y ~ Group | 1.79 | 2 | 0.41 |
| | Maximum heart rate | Gaussian | y ~ Group | 9.37 | 2 | 0.0092** |
| Novel object | Mean heart rate | Gaussian | y ~ Group | 2.87 | 2 | 0.29 |
| | Maximum heart rate | Gaussian | y ~ Group | 1.51 | 2 | 0.47 |
| All tests | Variation in cortisol level | Gaussian | Y ~ Group | 0.99 | 2 | 0.61 |

**Table S5: Descriptive statistics and effect sizes for the behavioral and physiological variables.** Effect sizes are given as the rate ratios (RR) or Cohen's d (d), as appropriate for each variable.

| Test | Response variable (y) | Condition | Mean±s.e. | Effect size (mean±se) | |
|---|---|---|---|---|---|
| Grooming | Number of touching the experimenter | Joy | 3.36±2.84 | | RR=1.18±0.28 |
| | | Neutral | 2.80±2.27 | | RR=2.79±0.21 |
| | | Fear | 1.43±1.40 | | RR=0.54±0.35 |
| Human approach | Number of touching the experimenter | Joy | 7.79±4.64 | | RR=1.11±0.20 |
| | | Neutral | 6.60±3.20 | | RR=6.83±0.15 |
| | | Fear | 4.14±2.93 | | RR=0.60±0.24 |
| | Mean heart rate | Joy | 58.64±10.16 | Joy-Neutral | d=-0.22±0.39 |
| | | Neutral | 61.67±15.12 | Neutral-Fear | d=-0.062±0.39 |
| | | Fear | 62.52±16.66 | Joy-Fear | d=-0.28±0.39 |
| | Maximum heart rate | Joy | 110.92±34.56 | Joy-Neutral | d=0.086±0.39 |
| | | Neutral | 107.62±43.23 | Neutral-Fear | d=-0.32±0.34 |
| | | Fear | 120.13±41.93 | Joy-Fear | d=-0.24±0.39 |
| Suddenness | Intensity of startle | Joy | 1.39±0.49 | Joy-Neutral | d=0.326±0.37 |
| | | Neutral | 1.23±0.42 | Neutral-Fear | d=-1.20±-0.40 |
| | | Fear | 1.82±0.61 | Joy-Fear | d=-0.88±0.39 |
| | Mean heart rate | Joy | 76.77±16.91 | Joy-Neutral | d=0.029±0.45 |
| | | Neutral | 76.36±12.63 | Neutral-Fear | d=-0.52±0.44 |
| | | Fear | 83.72±14.57 | Joy-Fear | d=-0.49±0.44 |
| | Maximum heart rate | Joy | 129.30±28.54 | Joy-Neutral | d=0.14±0.45 |
| | | Neutral | 125.03±20.55 | Neutral-Fear | d=-1.31±0.47 |
| | | Fear | 163.85±40.23 | Joy-Fear | d=1.16±0.47 |
| Novel object | Number of gazes | Joy | 5.21±2.49 | | RR=0.90±0.16 |
| | | Neutral | 5.80±2.96 | | RR=5.80±0.11 |
| | | Fear | 7.64±3.43 | | RR=1.32±0.14 |
| | Number of contacts | Joy | 2.07±2.62 | | RR=1.12±0.43 |
| | | Neutral | 1.53±1.73 | | RR=1.63±0.32 |
| | | Fear | 0.36±0.63 | | RR=0.32±0.58 |
| | Mean heart rate | Joy | 66.47±15.72 | Joy-Neutral | d=-0.73±0.45 |
| | | Neutral | 78.03±14.12 | Neutral-Fear | d=0.14±0.40 |
| | | Fear | 75.89±19.04 | Joy-Fear | d=-0.60±0.44 |
| | Maximum heart rate | Joy | 118.08±29.91 | Joy-Neutral | d=-0.50±0.45 |
| | | Neutral | 131.87±33.51 | Neutral-Fear | d=0.031±0.40 |
| | | Fear | 131.01±23.21 | Joy-Fear | d=-0.47±0.44 |
| PCA | F1 | Joy | 0.72±1.36 | Joy-Neutral | d=0.32±0.37 |
| | | Neutral | 0.38±1.21 | Neutral-Fear | d=1.40±0.40 |
| | | Fear | -1.13±0.68 | Joy-Fear | d=1.71±0.43 |
| | F2 | Joy | -0.077±1.43 | Joy-Neutral | d=-0.21±0.37 |
| | | Neutral | 0.15±1.04 | Neutral-Fear | d=0.22±0.37 |
| | | Fear | -0.089±0.92 | Joy-Fear | d=0.011±0.38 |
| All tests | Variation in cortisol level | Joy | 0.86±1.79 | Joy-Neutral | d=-0.20±0.27 |
| | | Neutral | 1.18±1.76 | Neutral-Fear | d=0.25±0.27 |
| | | Fear | 0.78±1.73 | Joy-Fear | d=0.052±0.27 |

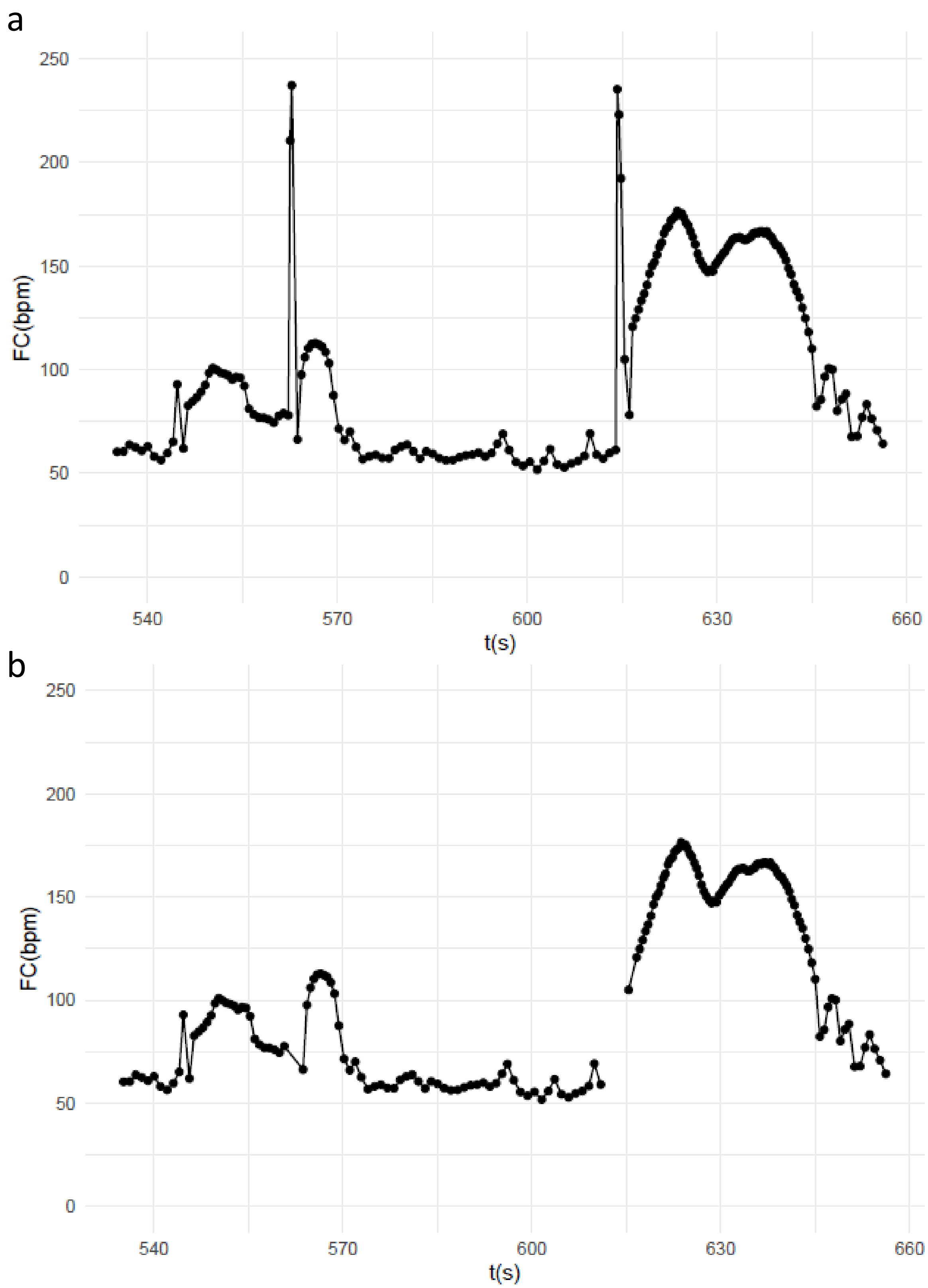


**<u>Figure S1</u>: Example of heart rate data before (a) and after (b) clearing up to cut artefactual beats.** The method used was examining each data point; if it had a difference of more than 35 bpm with the previous point, it was considered artefactual and removed from the data set and replaced by a missing value (NA).

CC BY

**Figure S2: Correlations between the behavioural variables**

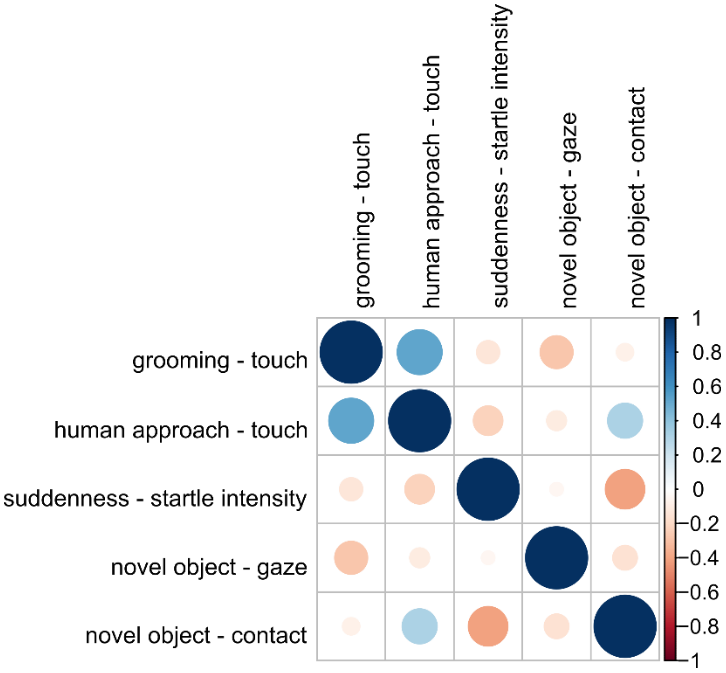
